# Phosphorylation inhibits intramolecular interactions, DNA-binding and protein interactions of Claspin through disordered/ structured conformation transition

**DOI:** 10.1101/2024.01.08.574761

**Authors:** Zhiying You, Hao-Wen Hsiao, Chi-Chun Yang, Hidemasa Goto, Hisao Masai

## Abstract

Claspin, known to be highly disordered, plays important roles in replication fork progression, initiation and cellular responses to replication stress. However, regulation of its structure and molecular interactions is not completely understood. We show here, through Proximity-Ligation-Assays, the evidence for intramolecular interaction between the N- and C-terminal segments of Claspin, which depends on the Acidic-Patch [AP] segment near its C-terminus. Interaction of Claspin with DNA and replication factors is highly stimulated in ΔAP mutant and by prior dephosphorylation. The wild-type Claspin inhibits the helicase activity of MCM in an AP-dependent manner. ΔAP and dephosphorylated Claspin exhibit resistance to trypsin digestion compared to wild-type, suggesting the presence of structural domains in the formers. We propose that Claspin is converted from disordered (closed) to structured (open) conformation at initiation, which stimulates its DNA binding and interaction with replication factors and counteracts its helicase inhibitory activity to trigger initiation of DNA replication.

## Introduction

At the eukaryotic DNA replication fork, the CMG (Cdc45-Mcm-GINS) complex serves as a replicative helicase, in which the Mcm hexameric complex plays a major role as an active unwinding enzyme. At the progressing replication fork, other accessory factors including Claspin, Tim, Tipin and And-1 play crucial role in facilitating the fork movement and stabilizing replication forks upon replication stress, generating a large replisome progression complex (RPC)^1,2,3,4,5,6^.

Claspin/Mrc1 is an evolutionally conserved replication checkpoint mediator and plays roles in the normal replication fork progression and in cellular response to stalled replication fork. It interacts with many replication factors including MCMs, Cdc7 kinase, Cdc45, PCNA, DNA polymerases α, 8, χ, ATR, Chk1, Tim, MCM10, and And-1^7,8,9,10^, consistent with its expected regulatory roles at the replisome. Indeed, Claspin is required for efficient fork progression in both yeast and human cells^9,11^, and yeast Mrc1 was shown to stimulate DNA replication through accelerating the unwinding rate of the replicative helicase in an in vitro reconstituted replication system, and this activity is regulated by phosphorylation^12,13,14,15^. Claspin may regulate the rates of DNA replication by coordinating template unwinding and leading-strand synthesis, ensuring replisome coupling to limit ssDNA exposure.

The fork protection complex (FPC), composed of the Claspin together with Timeless (Tim)-Tipin heterodimer (Tof1–Csm3) and And-1 scaffolds the replisome to ensure unperturbed fork progression and acts as a mediator of the DNA replication checkpoint at stalled forks^3,16^. In vitro, Claspin stimulates leading-strand replication with the help of Tim–Tipin^17^. Furthermore, Claspin facilitates initiation by recruiting Cdc7 through Claspin and promoting MCM phosphorylation, notably in non-cancer cells^10^. Cdc7 is a conserved kinase that triggers replication initiation through phosphorylating MCM. At the stalled replication fork, Claspin is required for the transduction of replication stress signals from ATR kinase (sensor kinase) to Chk1 kinase (effector kinase)^6,18^. Claspin recruits Chk1 to its Chk1 kinase binding domain (CKBD), which has been phosphorylated by Cdc7 or CK1λ1 to elicit successful checkpoint signaling induction^19,20,21,22,23^. Structurally, Claspin has an amino-acid compositional bias, with a low content of hydrophobic amino acids and a high content of polar and charged amino acids, which is typical for intrinsically disordered proteins. Indeed, a Cryo-EM structural study revealed that Claspin is highly disordered and only a few fractions of the polypeptides can be resolved^8^. Claspin may bind across one side of the replisome and contacts TIM and MCM subunits. Yeast Mrc1 was also reported to be associated with Tof1, Mcm6, Mcm2, and Cdc45^7^.

We have previously identified a domain (AP, acidic patch) that interacts specifically with Cdc7^10^. We suggested a possibility that AP may be involved in intramolecular interaction with its N-terminal segment. AP(DE/A) mutation of AP in which all the acidic residues are replaced by alanine results in loss of Cdc7 binding and that of the interaction with the N-terminal segment, leading to increased DNA and PCNA binding. However, effects of phosphorylation on the intramolecular interaction of Claspin and potential conformational transition of Claspin in the cellular context has not been examined. In the present study, we attempted to understand the effect of phosphorylation on the conformational transition and on its DNA and protein interactions as well as its functional significance. We found that the phosphorylation inhibits intramolecular interactions and DNA-binding activity of Claspin through significant conformational change. Proximity Ligation Assay (PLA) confirms that the interaction signal is strong with the wild-type Claspin but is very weak with AP(DE/A) or AP deletion mutants, demonstrating the intramolecular interaction of Claspin is dependent on AP. Furthermore, we discovered that Claspin can inhibit the helicase activity of Mcm in a manner dependent on the intramolecular interaction, suggesting a functional role of phosphorylation-induced structural transition. We propose that Claspin, highly disordered throughout the molecule, is converted at the initiation to the protein with structured domains, and is functionally transformed for replisome assembly.

## Results

### DNA-binding activity of Claspin is inhibited by phosphorylation

It has been known that Claspin preferentially binds to fork DNA^24,25^. Our laboratory previously reported that the AP (acidic patch) region of Claspin (aa 986–1,100), highly enriched in acidic residues (aspartic acid and glutamic acid), suppresses DNA-binding activity of the N-terminal segment of Claspin^10^. Using the purified polypeptides derived from Claspin, we conducted DNA-binding assays with a Y-fork DNA substrate. We found that the wild-type full-length as well as its derivatives, a degron-mutant and a double-tagged full-length, generated only smeared bands, suggesting that affinity is low or the complex, even formed, may be unstable during gel electrophoresis. On the other hand, AP(DE/A) mutant in which all the aspartic acid and glutamic acid residues in AP were substituted with alanine (aa 988–1,086) and ΔAP mutant in which the internal AP region was deleted generated distinct shifted bands (**Fig. 1A**). The purified Mcm2∼7 complex generated a shifted band on the Y-fork DNA, as previously reported^26^. All these Claspin-derived polypeptides generated additional slow migrating bands on the Y-fork DNA in the presence of the Mcm2∼7 (**Fig. 1A**, lanes 9-14 and lanes 18-19), similar to Cdt1^27^(**Fig. 1A**, lanes 15-16 and lanes 20-21). In the absence of DNA, the stable complex formation of Mcm2∼7 and Claspin is undetectable on a native gel (data not shown). The protein preparations used in this study were purified from mammalian cells, and may be already phosphorylated by endogenous kinases. We therefore dephosphorylated the affinity-purified Claspin protein with λPPase, and further purified the untreated and dephosphorylated preparations on a Mono Q chromatograph column. Both Claspin preparations eluted around 0.5 M KCl (data not shown). The untreated Claspin, presumably phosphorylated by endogenous kinases, exhibited only smeared bands as in Figure 1A, whereas λPPase treated preparation exhibited a distinct shifted band (**Fig. 1B**, lanes 2-3). On the other hand, both phosphorylated and dephosphorylated Claspin converted the Mcm2∼7-DNA complex to a slow-migrating form (i.e. the generation of the DNA-MCM-Claspin ternary complex) (**Fig. 1B**, lanes 5-6).

**Figure 1.**
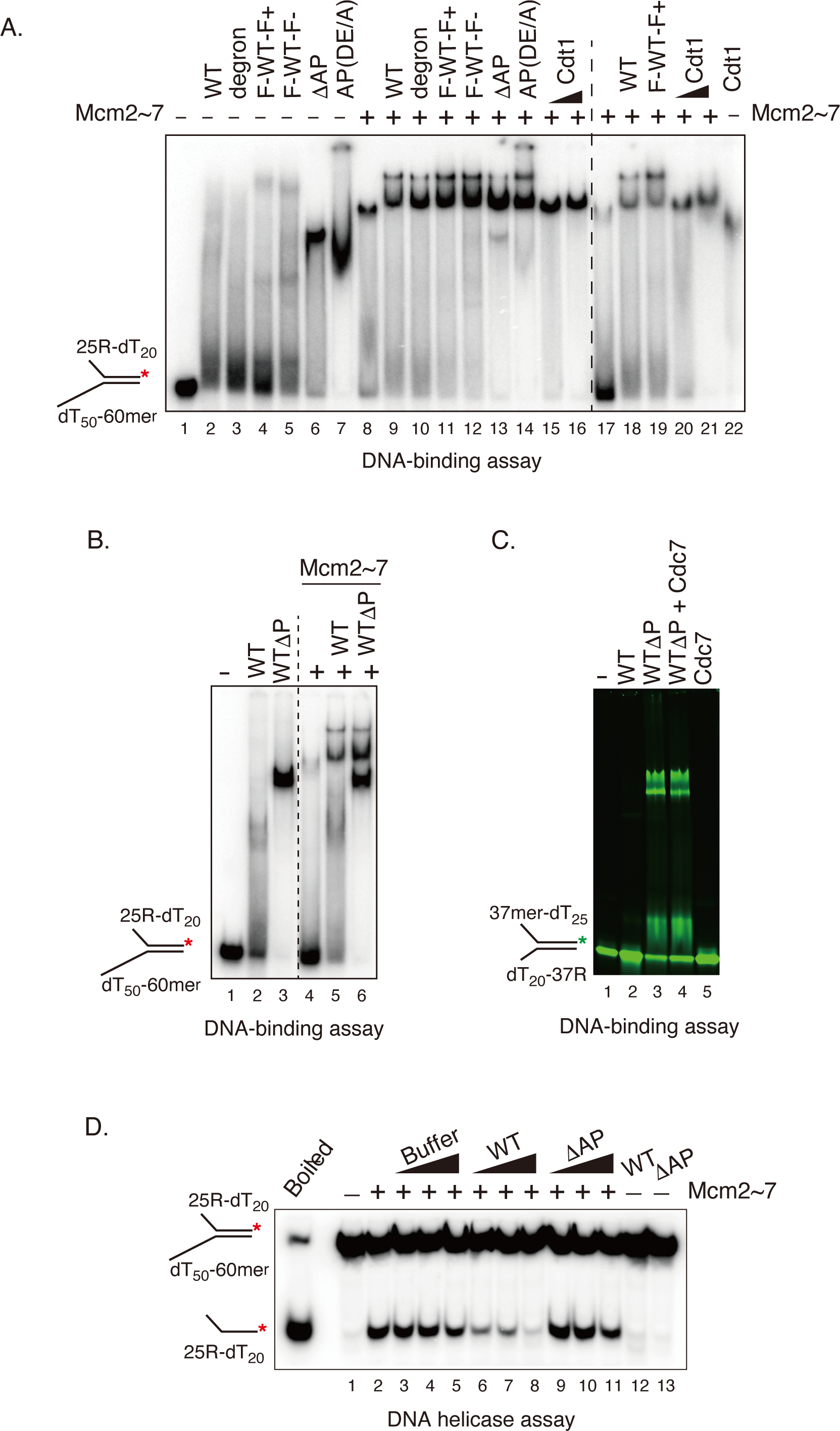
Phosphorylation of Claspin inhibits its DNA-binding activity. **A**. Gel shift assays of Claspin and its derivatives purified from 293T cells were conducted on Y-fork DNA. Purified proteins (300 ng each) used for assays were as follows; WT, full-length wild-type protein; degron, a mutant with N-terminal degron motif DS_30_GQGS_34_ replaced by DA_30_GQGA_34_; F-WT-F+ and F-WT-F-, FKBP and FRB tagged at the N- and C-terminus, respectively (+ or –, grown in the presence or absence of rapamycin); ΔAP and AP(DE/A) mutants. Proteins were added singly (lanes 1-7, 22) or in combination with the Mcm2∼7 complex proteins (lanes 8-21; 200 ng in leans 8-16 and 75 ng in lanes 17-21). As a positive control of complex formation, purified Cdt1 (50 and 100 ng) was used (lanes 15-16, 20-22). Y-fork DNA, 50 fmol. **B**. Dephosphorylation of Claspin activates DNA-binding of Claspin. 200 ng of untreated or λPPase-treated purified Claspin protein (lanes 2,3,5,6) and 75 ng of MCM protein (lanes 4-6) were used. Y-fork DNA, 50 fmol. **C**. Effect of Cdc7 on DNA-binding activity of Claspin using IR_800_-labeled Y-fork DNA. IR_800_-Y-fork DNA, 0.1 pmol; proteins added, 300 ng Claspin (untreated: WT or λPPase-treated: WT ΔP) and 50 ng Cdc7 protein. **D**. Effect of Claspin on DNA helicase activity of Mcm2∼7. Proteins added; highly purified Claspin protein (WT and ΔAP) 25, 50, and 100 ng; Mcm2∼7, 50 ng. Y-fork DNA, 15 fmol. As a control, the buffer of the same fraction of glycerol gradient fractionation was added (Buffer). Boiled, the Y-fork substrate after incubation at 96°C for 3 min.

We reported that Cdc7 interacts with the AP region of Claspin, and suggested that Cdc7-mediated phosphorylation of Claspin may reduce the intramolecular interaction between AP and N-terminal segment which may activate the DNA-binding activity^10^. To test this possibility, the purified dephosphorylated Claspin preparation, fully active in DNA binding, was phosphorylated by Cdc7 kinase with cold ATP and then the DNA-binding activity was examined. We found that Cdc7-mediated phosphorylation of Claspin did not affect DNA-binding of dephosphorylated Claspin (**Fig. 1C**). This is probably expected since Cdc7-mediated phosphorylation is predicted to activate DNA binding of Claspin. Claspin is phosphorylated in response to replication stress (HU addition). Therefore, we purified Claspin from cells (transiently transfected with a Claspin expression plasmid) treated with 2 mM HU for 1.5 hrs. DNA binding activities of both the WT and ΔAP proteins were not affected by pretreatment of the cells with HU either in the absence or presence of Mcm2∼7 (**Supplementary Fig. 1A**). Taken together, our results demonstrated that DNA-binding activity of Claspin is inhibited by phosphorylation, and the inhibition depends on the AP segment. Claspin forms a ternary complex with Mcm2∼7 on Y-fork, and this activity is not affected by the phosphorylation state of the Claspin.

### Claspin inhibits DNA helicase activity of the Mcm2∼7 hexamer

Next, we examined the effect of Claspin on Mcm2∼7 helicase activity. The wild-type Claspin was found to inhibit the Mcm2∼7 helicase activity in a dose-dependent manner compared to addition of buffer alone (**Fig. 1D**). In contrast, the ΔAP mutant did not significantly affect the Mcm helicase activity (**Fig. 1D**). In purification of WT Claspin through glycerol gradient centrifugation, the inhibitory activity on Mcm helicase was coeluted with the Claspin protein peak (data not shown). Both phosphorylated and dephosphorylated forms of Claspin exhibited similar inhibitory effect on the Mcm helicase (**Supplementary Fig. 1B**). Both dephosphorylated Claspin and ΔAP mutant have a high affinity to DNA and formed a ternary complex with Mcm2∼7, but only ΔAP lost the inhibitory activity, suggesting that the presence of AP rather than elevated DNA binding activity is responsible for Mcm helicase inhibition.

The recent reports indicate the physical interactions of Pol ε with Claspin, which may serve as a scaffold of replisome or as a regulator of catalysis^14^. Thus, we examined the effect of Claspin on DNA polymerase activity of Pol ε using the primer/template extension method. We found that stimulatory effect of Claspin on Pol ε did not coelute with Claspin (data not shown), indicating that Claspin does not stimulate the Pol ε activity, while *S. cerevisiae* Mrc1 was reported to stimulate the activity of DNA polymerase ε^28^.

### Phosphorylation reduces intra- and inter-molecular interaction by conformational change

Our previous studies demonstrated that the acidic patch (AP) near the C-terminus of Claspin can interact with N-terminus^10^. Claspin phosphorylation and expression are cell cycle-dependent, with Claspin levels undetectable in G1 cells, increasing in late G1/S phase, peaking in S phase, and decreasing in G2/M phase^29^. In accordance with the levels of Claspin during the cell cycle, the phosphorylation of Claspin increases during S phase. To examine whether this phosphorylation could be involved in regulation of intramolecular interaction between N- and C-terminal segments of Claspin, we first examined interaction between the N-terminal polypeptide (# 2; 1-350aa) and C-terminal polypeptide (#13 [897-1209aa] and #18 [897-1339aa], containing CKBD and AP), which were purified using the Flag and HA tags (**Fig. 2A**). To obtain dephosphorylated Claspin, we pretreated the lysate with λPPase. On the other hand, cells were treated with Calyculin A, which strongly inhibits protein phosphatase 1 (PP1) and protein phosphatase 2A (PP2A) that are responsible for the majority of protein dephosphorylation in cells, to obtain hyperphosphorylated Claspin. Claspin was also prepared from the cells in which Cdk1 or Cdk1/Cdk2 activity was inhibited by an inhibitor. #13 and #18 polypeptides were efficiently pulled down with Claspin #2 under dephosphorylation condition (pretreated by λPPase; **Fig. 2B, right**, lanes 5 and 6). In contrast, only a very small amount of #13 and #18 polypeptides was pulled down under hyper-phosphorylation condition (pretreated by Calyculin A; **Fig. 2B, right**, lanes 14 and 15). In non-treated or CDK inhibitor (RO-3306 and roscovitine)-treated cells, #13 and #18 polypeptides were pulled down at an intermediate level (**Fig. 2B, right**, lanes 2 and 3; lanes 8 and 9; lanes 11 and 12). Claspin would be phosphorylated by various kinases, and indeed, the N-terminal segment (#2) exhibited hyperphosphorylation after Calyculin A treatment (**Fig. 2B**, lanes 13-15). Thus, these results suggest that phosphorylation of Claspin at the N-terminal segment inhibits the intramolecular interaction between N- and C-terminal segments of Claspin. Furthermore, PCNA was also pulled down with #2 polypeptide under dephosphorylation condition, but not under hyperphosphorylation condition (**Fig. 2B**, compare lane 4 and 13). In addition, compared to #2, PCNA pull down was lost or reduced in #2+#13 and #2+#18 (**Figure 2A, right**, compare lanes 4, 7, and 10 with lanes 5,6,8,9,11 and 12), suggesting that intramolecular interaction of Claspin inhibits binding of PCNA to the PIP box (aa 311-318) of Claspin.

**Figure 2.**
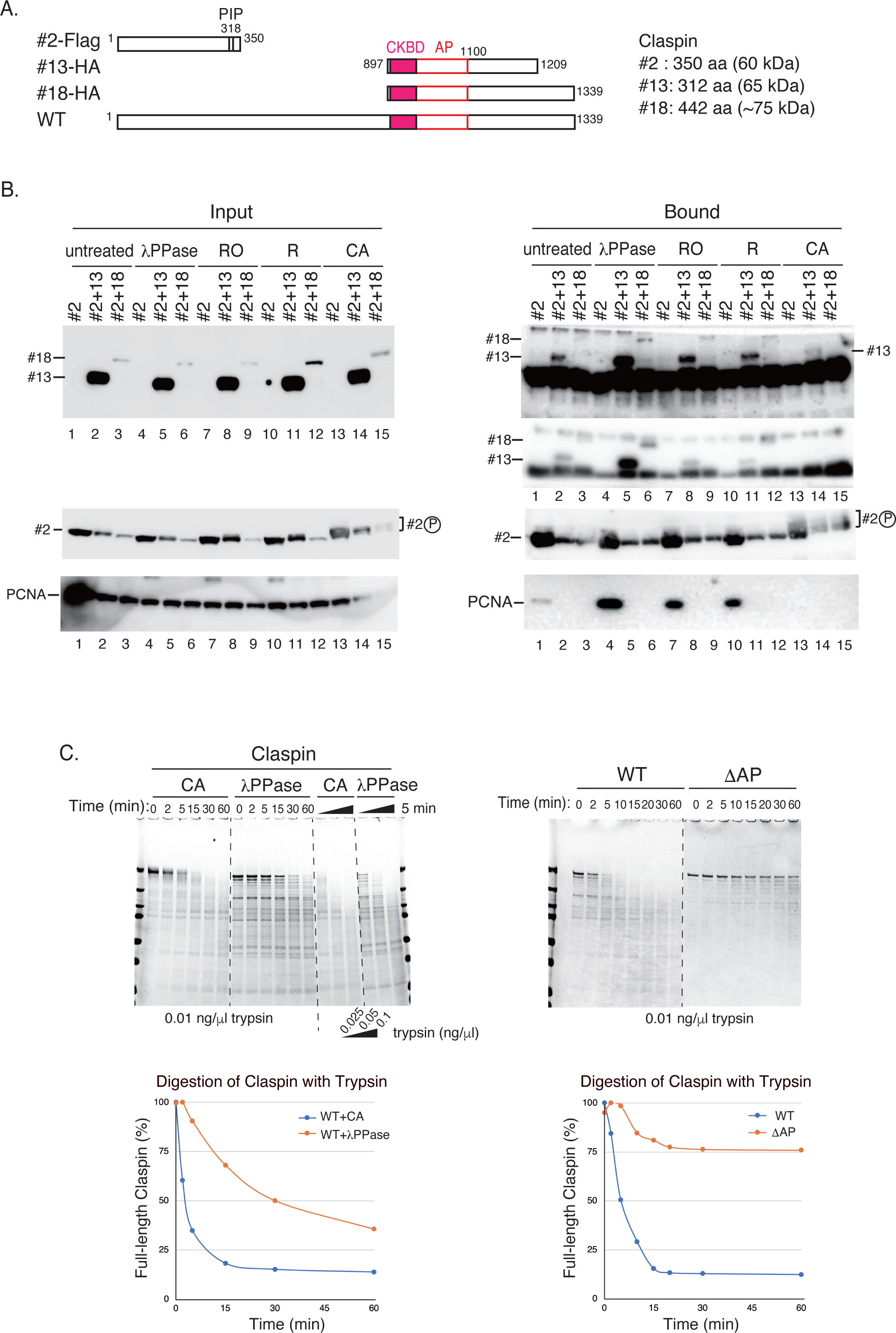
Interactions of the Claspin N-terminal segment with the C-terminal segment (CKBD+AP) and with other replication factors are regulated by phosphorylation. **A**. A schematic diagram of the various truncation mutants of Claspin used in the assay. At the C-terminus, #2 contains a 3x Flag and #13 and #18 carry a HA tag. **B**. Coimmunoprecipitation of N- and C-terminal polypeptides of Claspin. The Flag-tagged N-terminal polypeptide (#2) was coexpressed with the HA-tagged C-terminal polypeptides (#13 or #18). Cells were treated with 10 µM RO3306 (RO) or 20 µM Roscovitine (R) for 2 hr, or 100 nM Calyculin A (CA) for 30 min before cell harvest, where indicated. Cell lysates were lysed with native lysis buffer, and a half of non-treated cell lysates were incubated with λPPase. The inputs (Input) and immunoprecipitates with M2 anti-Flag antibody beads (Bound) were analyzed by western blotting using anti-HA, anti-Flag, and anti-PCNA antibodies. **C**. Proteolytic digestion of Claspin with trypsin. Purified Claspin treated with λPPase or Claspin purified from Calyculin A-treated cells was subjected to proteolytic digestion with 0.01 ng/µl trypsin (Sigma) in a time course, or with different concentrations of trypsin (0.025, 0.05, and 0.1ng/µl) in 25 mM Tris-HCl (pH 7.9), 50 mM KCl, 1 mM DTT and 0.1% Tween 20. In time course experiments, samples were incubated around 25°C and 12-µl samples were collected at indicated time points and quenched with SDS sample buffer. In enzyme titration, reaction was for 5 min. Samples were boiled and loaded onto an SDS–PAGE gel. Gels were stained with Coomassie Brilliant Blue (CBB) and band intensities of protein were measured using Multi Gauge software.

The above results suggest that the intramolecular interaction of Claspin as well as its interaction with other replication factors are regulated by phosphorylation. This may be caused by phosphorylation-induced conformation change of the Claspin molecule. We then digested phosphorylated or dephosphorylated Claspin with trypsin. Hyperphosphorylated Claspin was obtained from 293T cells treated with Calyculin A before cell harvest. Dephosphorylated Claspin was prepared from cell lysates pretreated with λPPase. The hyperphosphorylated Claspin migrated slower compared to the dephosphorylated form on SDS-PAGE (**Fig. 2C, left**). The phosphorylated form of Claspin is more sensitive to trypsin digestion than the dephosphorylated form both in the time course and enzyme titration experiments (**Fig. 2C, left**). The results suggest that phosphorylation changes the conformation of Claspin into a form less rigid (more vulnerable to attack of the protease) and less interactive. Consistent with this conclusion, ΔAP, which is expected to be underphosphorylated due to its inability to recruit kinases, was strongly resistant to trypsin digestion (**Fig. 2C, right**).

### Potential regulation of DNA binding by interaction between N-terminal and C-terminal domains

We next investigated the effect of intramolecular interaction on the ability of Claspin to interact with DNA. The truncated Claspin polypeptides including the N-terminal #2, C-terminal #13, #18, and #27 (containing CKBD and AP) were expressed in 293T cells and purified by using the anti-Flag antibody agarose (**Fig. 3A**). The dephosphorylated polypeptide would migrate more slowly in native gel than phosphorylated polypeptide, since negative charges are decreased by λPPase treatment. Indeed, an intrinsically N-terminal #2 polypeptide exhibited slow migration on the gel upon dephosphorylation (**Fig. 3C**, compare lanes 2 and 3). We examined DNA binding of these polypeptides using the IRDye 800CW near-infrared fluorescent dyes (IR_800_) labeled Y-fork DNA. The result shows that the #2 polypeptide binds to the DNA and the dephosphorylated form does so more efficiently than the phosphorylated form, whereas #13, #27 and #18 polypeptides (containing CKBD and AP) failed to bind to fork-DNA regardless of their phosphorylation state. This indicates that the N-terminal but not the C-terminal segment of Claspin has DNA binding activity (**Fig. 3B** and **3D, left**). In this assay, the wild type full-length Claspin binds to DNA efficiently only when it is dephosphorylated, as was shown in experiments using ^32^P labeled substrate (see **Fig. 1A**; **Fig. 3B, left**, lanes 14, 15 and **3D, left**, lanes 6 and 7).

**Figure 3.**
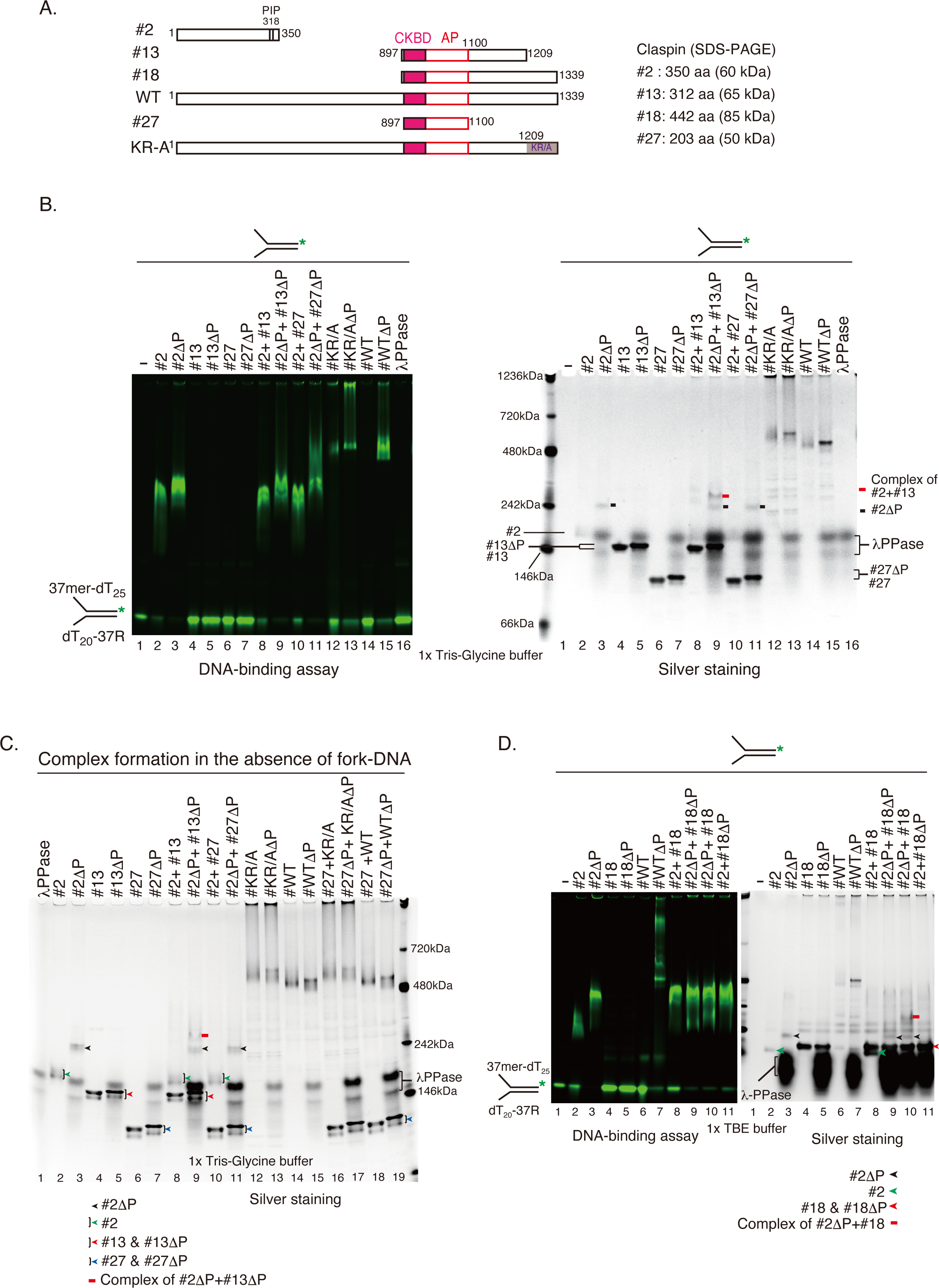
Interaction of N- and C-terminal segments of Claspin is observed in the absence of phosphorylation. **A**. A schematic diagram of the various truncation mutants of Claspin used in the assays. All the constructs contain a 3x Flag at the C-terminus. All the lysine and arginine residues in 1209-1339aa are replaced by alanine in the KR/A mutant. **B**. Gel-shift assays were conducted on IR_800_-labeled Y-fork DNA. Protein added, 300 ng; IR_800_ labeled Y-fork DNA, 0.1 pmol; λPPase, 100 units. Samples were run on a 5-20 % native gel. After scanning by Odyssey, the gel was stained by silver. **C**. Protein complex formation was examined in experiments similar to Figure 3B in the absence of DNA. **D**. Gel-shift assays were conducted as in B, except that proteins added were 200 ng and run on 1x TBE 5-20% native gel. After scanning by Odyssey, the gel was stained by silver. Proteins treated with 11PPase are labeled as ΔP.

We next examined the complex formation between the N-terminal and C-terminal polypeptides in the absence of DNA in native gel (**Fig. 3C**). Only the dephosphorylated #2 forms a complex with dephosphorylated Claspin #13 (**Fig. 3C**, lane 9, indicated by a red bar), consistent with the result of the pull-down assay in Fig. 2. Furthermore, in DNA-binding assay with Y-fork DNA, the proteins were detected with silver staining (**Fig. 3B, right**). The complex of #2 and #13 was detected only when dephosphorylated (**Fig. 3B, right**, lane 9, indicated by a red bar). These results indicate that the N-terminal #2 polypeptide forms a complex with C-terminal #13 polypeptide regardless of the presence or absence of DNA. Although the complex band was not detected in silver staining in combination of #2 and #27 treated with λPPase, smeared slow-migrating bands were detectable in more sensitive DNA-binding assay (**Fig. 3B, left**, lane 11). They strongly suggest a possibility of dephosphorylation-dependent intramolecular interaction of Claspin. Similar complex formation between #2 and #18 polypeptides was observed (**Fig. 3D, right,** lane 10, indicated by a red bar). In this experiment, wild type Claspin, # 2, and # 18 proteins were first treated by λPPase and the dephosphorylation activity was inactivated by phosSTOP, and then the gel-shift assays were conducted with IR_800_-labeled Y-fork DNA. The dephosphorylated #2 polypeptide formed a complex with both phosphorylated and dephosphorylated #18 polypeptide and Y-fork DNA (**Fig. 3D, left,** lanes 9 and 10). Since Claspin is rich in intrinsically disordered regions (IDRs) including 1-350aa segment of #2 polypeptide, phosphorylation can have a significant impact on protein folding and conformational change^30^. Indeed, dephosphorylated #2, unlike #13, #18, and #27, exhibited a significantly slower mobility (∼240 kDa) than untreated #2 (∼170 kDa) did in a native gel, which cannot be explained by changed charge (**Fig. 2**. B-D). The N-terminal segment #2 of Claspin (1-350aa) is predicted to be structured only at two segments (170-189aa and 277-299aa), while AP (988-1086aa) of Claspin is predicted to be unstructured. Furthermore, the complexes of dephosphorylated #2-#18-DNA migrated faster than dephosphorylated #2-DNA complex, showing that the ternary complex is more compact (**Fig. 3D**. left, lanes 3 and 9). Furthermore, the complexes #2-#13 or #2-#18 binds to DNA less efficiently than polypeptide #2 alone regardless the phosphorylation state from the remaining substrates (**Fig. 3B** and **3D, left**; and data not shown). The result suggests that the Claspin C-terminal # 13, # 18, and # 27 polypeptides containing CKBD and AP inhibit the DNA-binding activity of Claspin N-terminal region. This is consistent with the result that the Claspin derivative ΔAP lacking AP exhibit increased DNA-binding activity (**Fig. 1A**). Taken together, these results support the view that local phosphorylation or dephosphorylation reduces intramolecular interaction between N-terminal and C-terminal segments and activate DNA binding activity of the N-terminus segment.

### Claspin AP is essential for its intramolecular interaction as measured by PLA

The above results show the N-terminal and C-terminal segments can interact with each other. We next examined if this interaction contributes to the intramolecular interactions of Claspin in cells. For this, we have employed Duolink proximity ligation assay (PLA) that can visualize proteins present within the 40 nm proximity *in situ* (**Fig. 4A**). We constructed a derivative of Claspin which is tagged with HA or FKBP (FKBP12) and Flag at the N-terminus and the C-terminus, respectively, of Claspin. Using anti-HA or anti-FKBP and anti-Flag antibodies, red fluorescent spots (PLA signals) were detected in the wild-type Claspin, suggesting N-terminal (HA or FKBP) and C-terminal (Flag) segments interact through intramolecular interaction (**Fig. 4B** and **4C**). In contrast, the PLA signals were almost undetectable in the ΔAP mutant, suggesting that the intramolecular interactions requires AP. Similarly, PLA signals were not detected in AP(DE/A) mutant in which all the acidic residues in AP were replaced with alanine (**Fig. 4B**). These results strongly support the existence of an intramolecular interaction in Claspin that depends on the acidic residues in AP.

**Figure 4.**
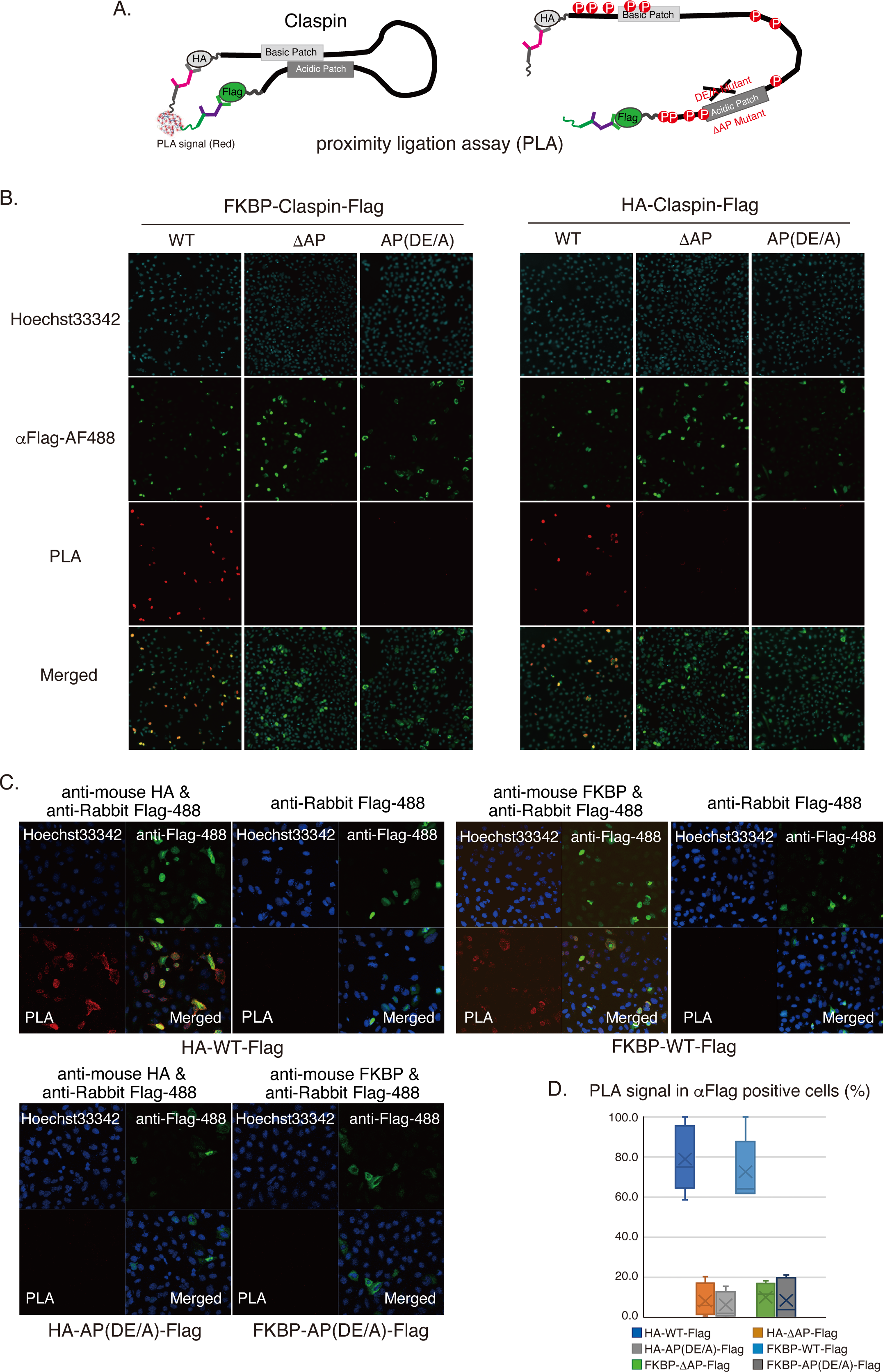
An essential role of AP in intra-molecular interaction of Claspin. **A**. Diagram of PLA using Claspin carrying HA and Flag tags at the N-terminus and C-terminus of Claspin, respectively. **B.** and **C**. PLA with FKBP-Claspin-Flag and HA-Claspin-Flag. Claspin: WT (wild-type), ΔAP, and AP(DE/A). Signals were observed by All-in-One fluorescence microscope (x10 expansion) (**B**) or confocal microscopy (**C**). Red, PLA signal; Green, Claspin (Alexa Fluor 488 anti-Flag antibody); blue, nuclei. **D**. Summary of PLA analysis. The graph shows the quantification of the PLA signals by calculating the ratio of PLA signals to anti-Flag signals. The data from 200 cells was statistically analyzed and a box plot has been used.

### N-C interaction in intramolecular interaction masks the N-terminal phosphorylation and exposes the C-terminal phosphorylation sites

DISPHOS (DISorder-enhanced PHOSphorylation precision) predicts that 128 of Claspin’s 201 serine and threonine residues are predicted to be phosphorylated. Phosphorylation of CKBD is required for Chk1 recruitment and activation. It was reported that casein kinase (CK1λ1) can phosphorylate CKBD and activate Chk1^20^. We reported that Cdc7, in conjunction with CK1λ1, can phosphorylate CKBD, and that Cdc7 play a predominant role in cancer cells, whereas CK1λ1 plays a major role in normal cells^19^. Furthermore, casein kinase (CK2) interacts with the replisome containing Claspin and may regulate the checkpoint activation^31^. To understand how phosphorylation affects the structure and functions of Claspin, the effects of various kinases including Cdc7, CK1λ1, Chk1, and CK2 were examined.

The kinase assays were performed using various Claspin derivatives including #2 (N-ter), #13 (C-ter) and #27 (C-ter) polypeptides, which were treated with λPPase and then with phosSTOP prior to addition to the kinase assay. We found that the polypeptide #2 was efficiently phosphorylated by all four kinases including Cdc7, Chk1, CK1λ1 and CK2. In contrast, #13 and #27 were phosphorylated by CK2 but not by CK1λ1, Chk1, or Cdc7 (**Fig. 5 A, B, and C**). The levels of phosphorylation of ΔAP lacking AP are reduced compared to those of the wild-type Claspin for all kinases (**Fig. 5 A, B, and C**, compare WT to ΔAP). This is probably due to the loss of recruitment of the kinases by ΔAP. #13 and #27 are not efficiently phosphorylated by CK1λ1, Chk1, or Cdc7, although they contain AP. This indicates that the phosphorylation sites on Claspin by these kinases lie not in the C-terminal segment, but in the N-terminal segment. Indeed, the N-terminal polypeptide #2 is efficiently phosphorylated by them. On the other hand, CK2 can phosphorylate #13 and #17, suggesting the presence of CK2-mediated phosphorylation sites in the C-terminal segment (**Fig. 5A, B, and C**).

**Figure 5.**
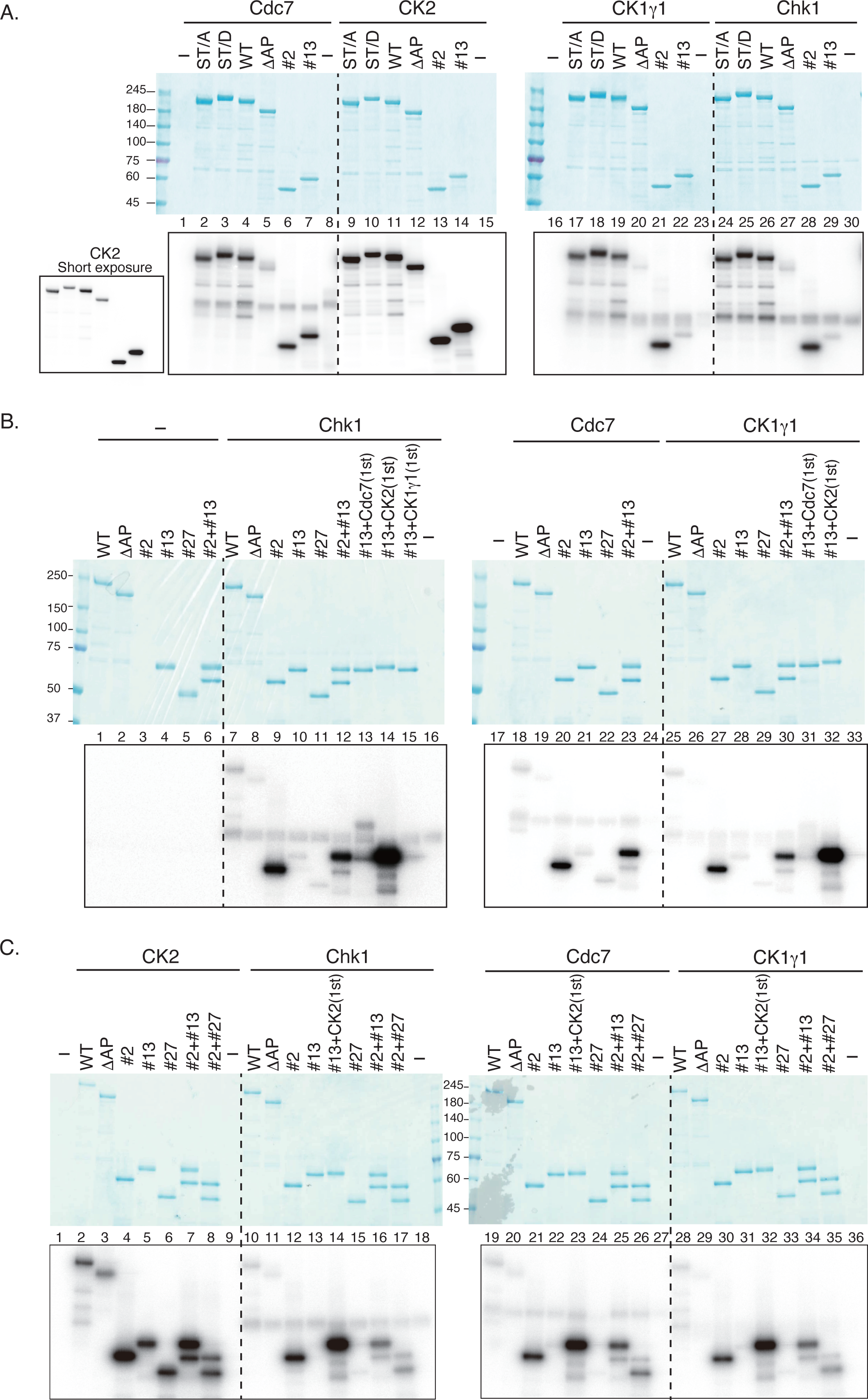
Claspin is phosphorylated by various kinases. **A**. Kinase assays were conducted with various kinases (25 ng; CK2, 50 ng) indicated. Substrates used were 300 ng each of WT (wild-type), ST/A, ST/D, ΔAP, #2 (N-terminal), #13 (C-terminal) and #27 (C-terminal). Purified proteins were treated with λPPase and reactions were terminated by addition of phosSTOP. Unlike the results shown in Fig. 5 **B and C**, Cdc7 appears to phosphorylate #13 to some extent in this particular experiment for unknown reason (lane7). **B**. Kinase assays were conducted with a single kinase or combinations of kinases (25 ng each) indicated. **C**. Kinases assays were conducted with kinases (25 ng) and substrate polypeptides (300 ng) indicated. After kinase reaction, 5x sample buffer was added to the reaction mixtures, and the samples were heated at 96°C for 2 min and then applied to SDS-PAGE. After CBB staining, the gel was dried and autoradiographed. In the reactions with two kinases in B and C, the #13 polypeptide was incubated with indicated kinases (1^st^) in the presence of cold 50 µM ATP for 30 min at 30°C, followed by incubation with the 2^nd^ kinase (indicated above the bars) in the presence of γ-^32^P ATP. “#2+#13” and “#2+#27” indicate the presence of two substrate polypeptides.

We next examined the effect of intramolecular interaction on the phosphorylation of Claspin. For that, we mixed N-terminal #2 and C-terminal #13 or #27 and phosphorylated them by CK1λ1, Chk1, or Cdc7. As stated above, #2 polypeptide is a good substrate of these kinases (**Fig. 5B**), but its phosphorylation level is significantly reduced, and that of the coexisting C-terminal polypeptides is significantly increased, when the two polypeptides were mixed (**Fig 5B**, lanes 12, 23 and 30; **Fig. 5C**, lanes 16-17, 25-26, and 34-35). This occurs with three different kinases. We speculate that the N-terminal phosphorylation sites are masked when #2 is complexed with the C-terminal polypeptides. At the same time, phosphorylation sites on the C-terminal polypeptides are exposed, leading to their phosphorylation. Additionally, the complex formation may facilitate the interaction of the kinases (specifically Cdc7 and CK1λ1, known to be acidophilic kinases) with AP. This suggests that these kinases can phosphorylate both N-terminal and C-terminal segments of Claspin in the “closed” conformation.

We next investigated the combination of two kinases, since prior phosphorylation by a “priming” kinase could facilitate the “secondary” phosphorylation by another enzyme. Therefore, we mixed CK1λ1 with Cdc7 or CK2 and examined the phosphorylation of the C-terminal #13 polypeptide. We did not see any synergistic effect between CK1λ1 and Cdc7 or Chk1 (**Fig. 5B**, lanes 31 and 15), concluding that these kinases independently act on the C-terminal segment of Claspin. On the other hand, we observed increased phosphorylation of #13 in the presence of both Cdc7 and Chk1 (**Fig. 5B**, compare lane 10 to lane 13), while either kinase alone shows very little phosphorylation (**Fig. 5B**, lanes 10 and 21). This effect was not seen with the CK1λ1-Chk1 combination. This is probably due to the phosphorylation of CKBD by Cdc7 which facilitates the recruitment of Chk1. This speculation is supported by our previous results that Cdc7 is more active than CK1λ1 as a CKBD-phosphorylation kinase in vitro^19^.

The presence of both CK2 and Cdc7, Chk1, or CK1λ1 increased the phosphorylation of #13 compared to each kinase alone (**Fig. 5C**, compare lanes 5,13,14, 22,23, 31and 32). This suggests the synergy between CK2 and the three kinases for phosphorylating the C-terminal polypeptide. The physiological roles of this synergy or the role of CK2 are currently unknown.

### N-terminal segment is responsible for DNA binding of Claspin and is regulated by phosphorylation

The N-terminal polypeptide #2 appears to be highly phosphorylated, as indicated by mobility-shift upon phosphatase treatment (**Fig. 3B**, lanes 2 and 3). The same polypeptide can be efficiently phosphorylated in vitro by various kinases (**Fig. 5**). Therefore, we have generated mutants in which all serine and threonine in the N-terminal 350aa region were replaced with alanine (ST/A mutation) or phospho-mimic aspartic acid (ST/D mutation) on the full-length Claspin. After purification of these Claspin mutants in 293T cells, the effects of the mutations on the phosphorylation by kinases and on DNA-binding activity were investigated. All of them are phosphorylated by Cdc7, CK2, CK1λ1, and Chk1 kinases to similar levels (**Fig. 5A**, lanes 2-4; 9-11, 17-19, 24-26). Thus, these results suggest that N-terminal serine/ threonine mutations did not appreciably change the total phosphorylation level of Claspin in vitro by these kinases.

To investigate whether DNA-binding activity is affected by the ST/A and ST/D mutations, gel-shift assays were conducted using IR_800_-labeled Y-fork DNA. It should be noted that Claspin is highly phosphorylated when expressed in the 293T cells. In fact, the treatment with λPPase caused the gel mobility of the wild-type to become slower in native gel, suggesting that λPPase had removed negatively charged phosphate groups of Claspin (**Fig. 6A, right,** lanes 2 and 3; **Fig. 6B, right,** lanes 3-5). On the other hand, the ST/A mutant, in which the phosphorylation sites are replaced by alanine with uncharged side chains, migrated slower than the WT. In contrast, the ST/D mutant containing negatively charged aspartic acid (D) at N-terminus, migrated faster than the WT (**Fig. 6, right**). In gel-shift assays using Y-fork DNA, both WT and ST/A exhibited significant DNA binding only after dephosphorylation treatment (**Fig. 6A**). In contrast, the phosphorylation mimic ST/D showed no DNA-binding activity even after treatment of λPPase, consistent with previous results that the phosphorylation inhibits DNA-binding activity (**Fig. 6A**). It was unexpected that the ST/A mutant is inefficient in DNA binding. However, titration of protein amounts indicated higher DNA binding activity of ST/A than the WT (**Fig. 6B**, compare lanes 3-4 and lanes 7-8). This is consistent with the fact that DNA binding of the N-terminal polypeptide #2 increases upon dephosphorylation. The result also indicates that the major DNA binding activity of Claspin is mediated by the N-terminal segment.

**Figure 6.**
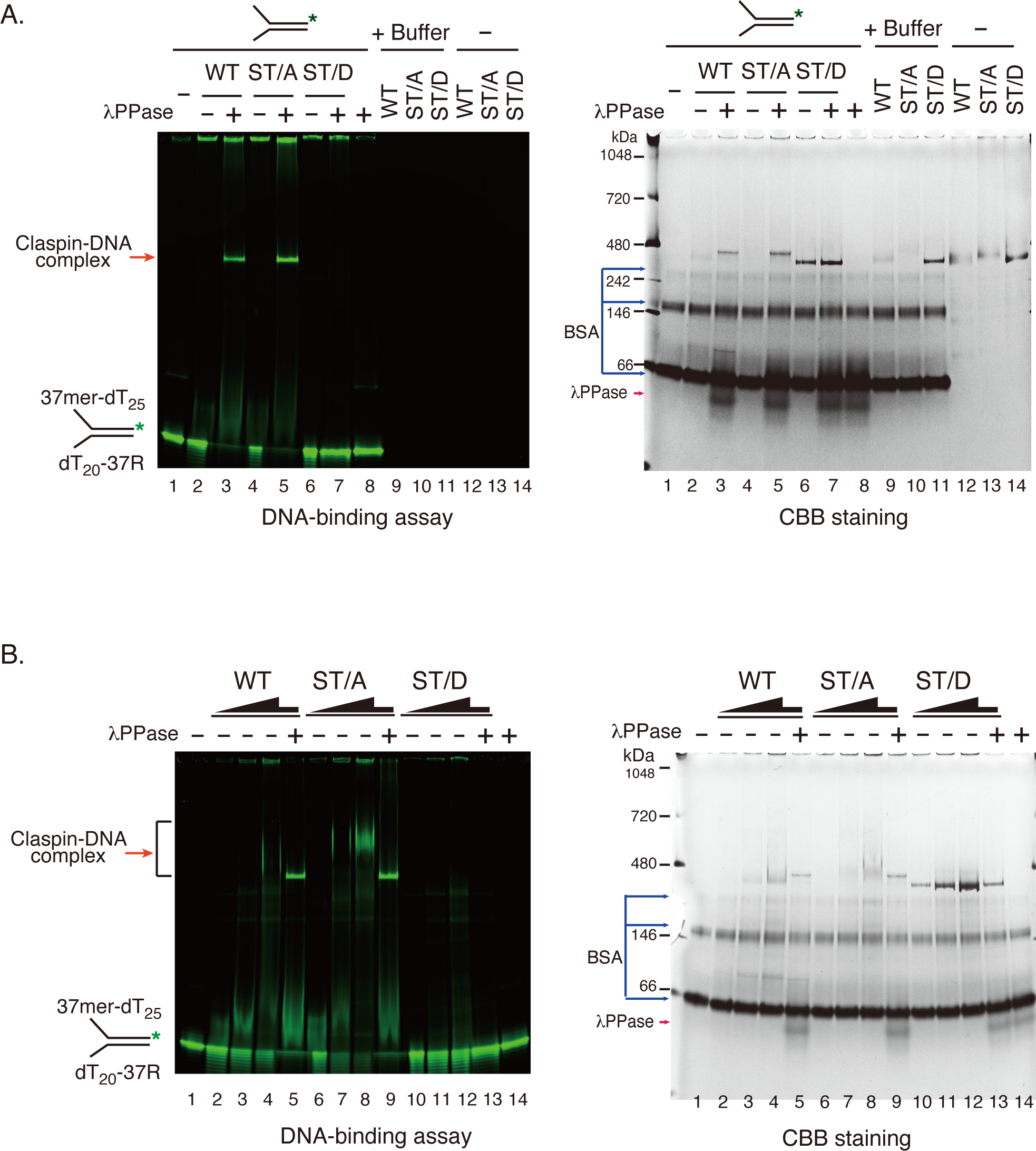
The Claspin N-terminal segment is responsible for binding to DNA, and is regulated by phosphorylation. **A.** and **B.** Gel-shift assays were conducted on IR800-labeled Y-fork DNA. Proteins used, WT (wild-type), ST/A, and ST/D mutants (300 ng in A, 300 ng, 600 ng, and 900 ng in B). +, treated with λPPase (100 units), -, untreated. Y-fork DNA, 0.1 pmol. Samples were run on 1x TBE 5-20% native gel. After scanning by Odyssey (left panel), the gel was stained by CBB (right panel). The reaction buffer contains BSA, which is indicated by blue arrows.

### Claspin ST/D mutant affects checkpoint activation in response to DNA damage

In view of the role of the N-terminal segment in DNA binding and its regulation by phosphorylation, we examine the functions of these mutants in the checkpoint response. For that, we have established HCT116 cell lines containing endogenous Claspin protein tagged with a minimum auxin-inducible degron (mAID) through CRISPR/Cas9-based gene editing. Thus, in this cell line, Claspin can be rapidly and specifically degraded in an indole-3-acetic acid (IAA; a natural auxin)-dependent manner. To ectopically express various Claspin derivatives, the relevant cell lines were constructed using lentivirus infection. The isolated lentiviruses were used to transfect human Claspin-mAID HCT116 cells and to establish the cell lines expressing the WT, ST/A, and ST/D Claspin. The addition of IAA induced depletion of endogenous Claspin-mAID, which was confirmed by western-blotting (**Fig. 7A**, compare Vector –IAA with +IAA; lanes 1 and 2).

**Figure 7.**
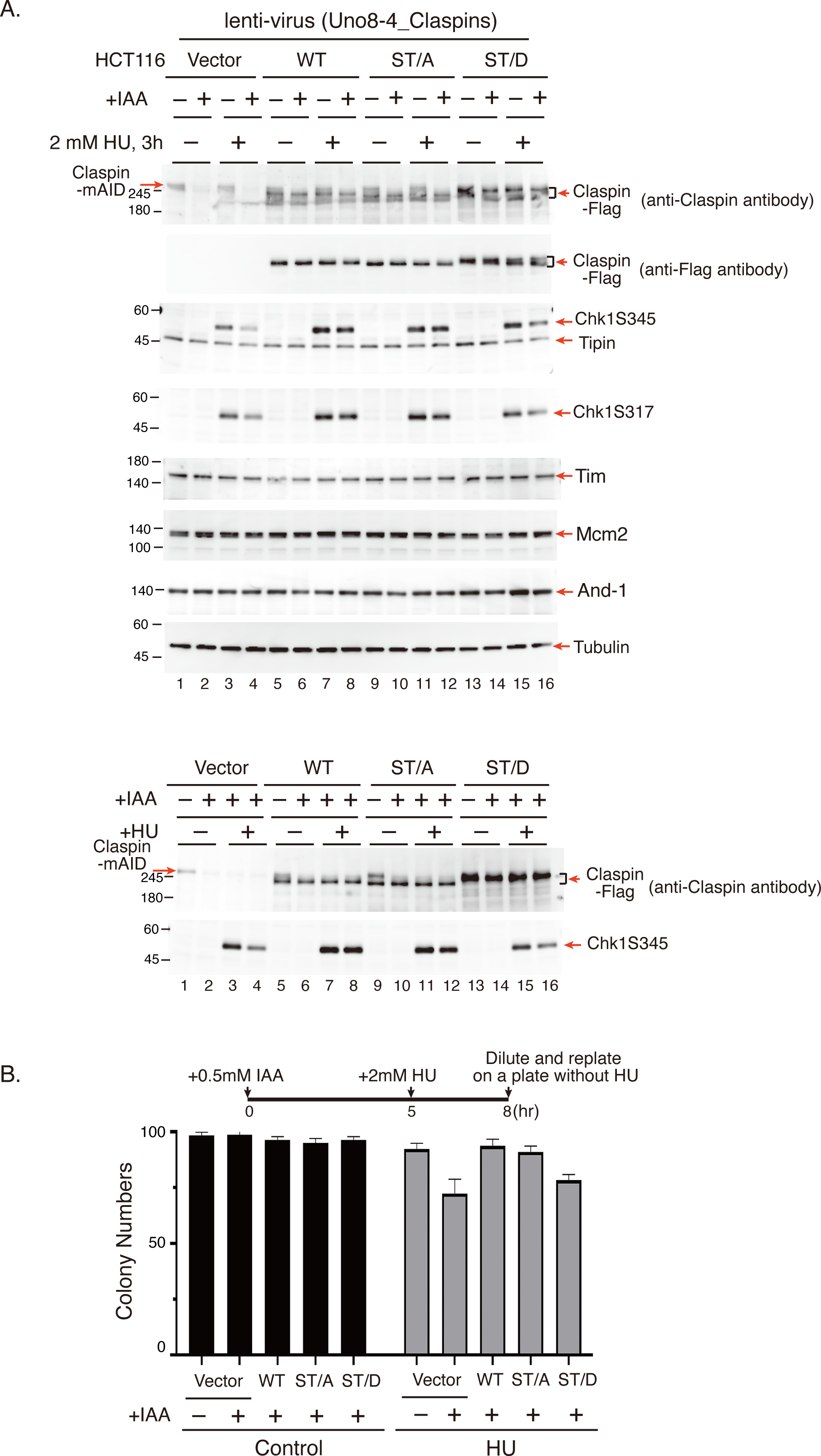
Claspin phosphomimetic ST/D mutant exhibits defective replication stress responses and poorer cell survival under stresses. **A.** The HCT116 Claspin-mAID cells expressing wild-type (WT) or mutant forms (ST/A and ST/D) of Claspin were first treated with IAA for 4 hr or untreated, as shown. Cells were then untreated or treated with 2 mM HU for 3 hr (upper) or 2 mM HU for 2 hr (lanes 3,7,11,15) and 10 mM HU for 1 hr (lanes 4,8,12,16) (lower), and were collected for western blot analysis of various proteins as indicated. **B.** The HCT116 Claspin-mAID cells expressing the wild-type or mutant Claspin were first treated with IAA for 5 hr as shown. Cells were then treated with 2 mM HU for 3 hr, and were replated onto a plate that did not contain HU after dilution. Cell survival rates of the WT, ST/A and ST/D cells, non-treated or treated with HU, were assessed by counting the numbers of the colonies formed and are shown in the bar graph. Error bars represent the mean values ± SEM under two independent experiments, both of which include three replicates.

In response to replication stress, Chk1 is phosphorylated at Ser317 and Ser345 which requires Claspin as a mediator (**Fig. 7A**, in the absence of IAA; compare lanes 1 and 3). On the other hand, IAA incubation reduced phosphorylation of Chk1 due to degradation of Claspin in the Claspin–mAID cell line, consistent with the requirement of Claspin for checkpoint activation (**Fig. 7A**, compare lanes 3 and 4). By ectopically expressing wild-type, ST/A, or ST/D mutant version of Flag-tagged Claspin (**Fig. 7A**, second panel from the top), the rescue experiment was conducted in the presence of HU. In the presence or absence of IAA, the wild-type and ST/A mutant showed similar levels of Chk1 phosphorylation (**Fig. 7A**, compare lanes 7 to lane 8, and lanes 11 to lane 12), indicating that ST/A mutant can rescue Chk1 activation in Claspin-depleted cells. On the other hand, in the ST/D mutant cells, Chk1-S345 and -S317 levels were similar to the vector control (**Fig. 7A**, compare lanes 15 and 16). These results indicate that the N-terminal phospho-mimic ST/D mutant is defective in checkpoint activation. Expression of other fork factors such as Tim-Tipin, And-1 and Mcm2 remains unchanged (**Fig. 7A**). Similar results were obtained in other HU treatment conditions (**Fig. 7A**, lower; 2 mM for 2 hr and 10 mM for 1 hr) and HU time-course experiments (10 mM, 1hr; and 2 mM, 2 and 3 hr; **Supplementary Fig. 2A**). We also conducted the colony formation assays to evaluate the effects of these mutation on cell proliferation after HU treatment. In cells treated with IAA and HU, colony numbers were reduced in the ST/D mutant similar to the vector control (**Fig. 7B**). These results show defective phenotypes of ST/D mutant cells in replication checkpoint activation and cell survival after replication stress.

Finally, to assess whether N-terminal phosphorylation affects the intramolecular interaction between Claspin N-terminal and C-terminal segments, we used HCT116 cell line containing endogenous Claspin-mAID and ectopically expressing the WT, ST/A or ST/D Claspin tagged with HA at the N-terminus and with Flag at the C-terminus. After Claspin-mAID depletion by addition of IAA, PLA signals could be observed not only in the wild-type Claspin cells, but also in ST/A and ST/D Claspin cells (**Supplementary Fig. 2B**). These results suggest that conformational change induced by the N-terminal phosphorylation, if any, may not cause the proximity change larger than 40 nm.

## Discussion

Intrinsically disordered regions (IDRs) are characterized by their lack of defined structure. Claspin/ Mrc1 is rich in IDR and only a small portion of the entire protein is structurally resolved by Cryo-EM by analyses of purified Claspin/ Mrc1 proteins which are presumably already phosphorylated since it was purified from yeast cells^7,8^. We found that the phosphorylation of Claspin inhibits its intramolecular interactions and DNA-binding and protein interacting activities presumably through conformational change of Claspin.

### Phosphorylation regulates the DNA-binding activity of Claspin through its effect on the intra-molecular interaction

We discovered that when Claspin was dephosphorylated, the DNA-binding activity is restored to the same level that is exhibited by ΔAP and AP(DE/A) **(Fig. 1**). Therefore, we speculate that specific, local phosphorylation can contribute to the closed conformation during G1. The N-terminal segment of Claspin is known to be required for DNA-binding in Xenopus and human^10,24,32^. Indeed, only the N-terminal segment #2 (1-350aa), but not C-terminal polypeptides, binds to DNA. The dephosphorylated form of the #2 polypeptide has much stronger DNA binding activity than the phosphorylated form (**Fig. 3**). Moreover, the ST/D mutant, in which N-terminal #2 segment of the full-length Claspin mimics hyperphosphorylation, completely loses its DNA binding ability, consistent with above results, whereas the ST/A mutant did not show a dramatic increase in DNA-binding over WT (**Fig. 6**). The DNA binding of the full-length Claspin, with the non-phosphorylated state of N-terminal segment, may be masked by its inhibitory interaction with the C-terminal segment. DNA binding activity of Claspin is regulated by both phosphorylation of the N-terminal DNA binding domain and by the intra-molecular interaction between the N -and C-terminal segments. This may be similar to p53, where its phosphorylation modulates the internal interactions of the IDR with the DNA-binding domain, which affects DNA binding activity^33^.

The phosphomimicking mutation, ST/D, in the N-terminal 350aa polypeptide impaired DNA binding and showed defect in replication check point responses. It may destabilize or weaken intramolecular interaction, although PLA signal is still detectable under our assay conditions (**Supplementary Fig. 2**). The reduced interaction between N- and C-terminal segments would lead to inefficient phosphorylation of the C-terminal segment including CKBD, which is essential for Chk1 activation. These results suggest that the N-terminal segment can be involved in regulated phosphorylation of Claspin on CKBD during checkpoint response.

### Cell cycle-dependent phosphorylation may trigger conformational change of Claspin

Claspin is a cell cycle-regulated protein the expression of which peaks at S/G2 phase^34,35,36^. During unperturbed growth, Claspin/Mrc1 is highly phosphorylated during late S to G2/M phase and underphosphorylated from G1 to early S phase^29,37^. Since Claspin is very rich in IDRs which are preferred targets of phosphorylation, it can be phosphorylated at multiple sites. Indeed, Claspin can be phosphorylated by Cdc7, CK1λ1, ATR, and Chk1, and also by other kinases, such as Polo-like kinase 1, stress-activated protein kinase (SAPK) p38 and CK2α^3,31^. In this report, we have shown that Claspin is directly phosphorylated by Cdc7, Ck1λ1, Chk1, and CK2α kinases in vitro (**Fig. 5**). Among them, CK2 is implicated in the phosphorylation of repair proteins, which are platform proteins for the recruitment of other proteins to the damaged DNA for repair^38^. It was reported that CK2α interacts with Claspin and CMG, and perturbation of the CK2α level results in defective S-phase checkpoint activation^31^.

The fork protection complex (FPC), composed of Claspin and Tim-Tipin, is essential for rapid and efficient replisome progression^9,11,12,39,40^. Structural studies show that Claspin has an extended and flexible configuration spanning one side of the replisome, from its N-terminal association with TIM, across MCM6 and MCM2, towards CDC45^7,8^. Reconstituted yeast replisomes indicate that Mrc1 can enhance fork rate and the phosphorylated Mrc1 is compromised in its activity^12,13^.

We have discovered that Claspin can inhibit DNA helicase activity of the Mcm hexamer. This activity depends on AP and prior dephosphorylation increased the inhibitory effect (**Fig. 1D**, **Supplementary Fig. 1B**). Although the helicase-inhibitory effect needs to be examined using the CMG helicase as well as using more functional assay systems (e.g. in vitro reconstituted replication system), our data suggests the presence of a novel mode of regulation of initiation by Claspin. The results also indicate that “closed conformation” is capable of inhibiting Mcm, while “open conformation” represented by ΔAP, is not. During the normal course of DNA replication initiation, Claspin may send a stop signal for the helicase, which would be released by Cdc7-mediated phosphorylation of Claspin, converting it to an “open conformation”. This is consistent with the finding in fission yeast that Mrc1 lacking HBS (Hsk1 binding sequence; Hsk1, fission yeast homologue of Cdc7 kinase) can bypass Cdc7 function presumably by removal of the inhibitory function of Mrc1^37,41^. Claspin constitutively in an open conformation may prematurely activate the origin in the absence of Cdc7 kinase^42^. During normal course of initiation, Claspin may transiently inhibit the unwinding and conversion to an open conformation may trigger initiation by counteracting the inhibition. Thus, loss of Claspin could lead to accelerated initiation of DNA replication. It has been reported that Claspin inactivation could be an essential event during carcinogenesis, suggesting a possibility that Claspin may function as a tumor suppressor^43^. In conclusion, we speculate that Claspin may play an important role at the initiation by changing its conformation from “closed” to “open”, which increases the affinity to DNA and replisome proteins and may also stimulate the helicase activity of the CMG complex^44^. It would be necessary to examine the cell cycle-dependent conformational change of Claspin by PLA in the future.

### Regulation of Claspin functions by potential conversion between disordered and structured conformations

It has been known that protein phosphorylation predominantly occurs within the IDR, and that multisite phosphorylation of IDR would have diverse effects on structure, conformational ensemble, and molecular functions^30,45^. On the basis of the data presented, we would like to propose an emerging model on how the phosphorylation state of Claspin affects the intramolecular interaction, overall structures and affinity to DNA and replisome proteins as well as the inhibitory activity on the unwinding mediated by Mcm2∼7 helicase (**Fig. 8**).

**Figure 8.**
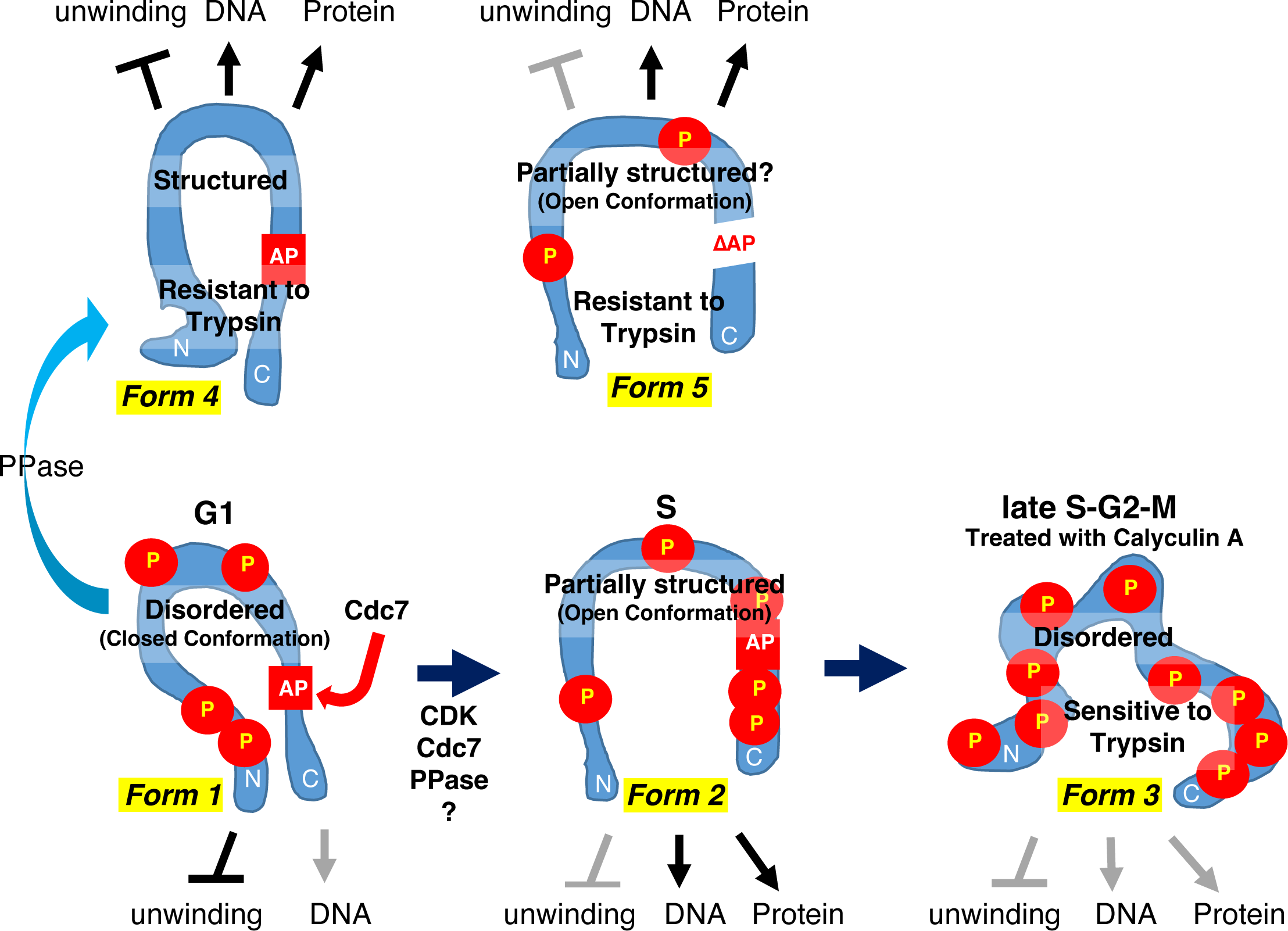
Summary of predicted structures and functions of Claspin during cell cycle. Claspin is depicted as a blue tube. P in circles represent phosphorylation. The black arrows indicate DNA binding and protein binding. The unwinding inhibition is shown by T-shaped bars. Black and grey indicate “strong” and “weak”, respectively. During G1, Claspin may be phosphorylated at a basic level, generating a closed conformation (Form 1), whose DNA binding activity may be masked by N-C interaction. This form may inhibit the helicase activity of Mcm. Upon entry into S phase, Claspin is asymmetrically phosphorylated or dephosphorylated by Cdc7, Cdk or a phosphatase and is converted to an open conformation (Form 2), which permits binding to DNA and other factors. This form may not inhibit the Mcm helicase any more. At late S-G2-M, Claspin may be hyper-phosphorylated and become highly disordered (Form 3; predicted from data in **Supplementary Fig. 3**), disabling its DNA and protein binding activities. This form can be induced by treatment with Calyculin A. The protein phosphatase treatment (light blue arrow) would convert Claspin into a highly structured form with DNA and protein interacting abilities (Form 4; predicted from data in **Supplementary Fig. 3**). Similarly, ΔAP mutant of Claspin lacks critical phosphorylation due to its inability to recruit kinases, and adopts a structured conformation (Form 5). Both Form 4 and Form 5 are resistant to Trypsin digestion, while Form 3 is highly sensitive, showing the correlation between trypsin sensitivity and IDR (non-structured)/ structured conformation transition.

Almost entire segment of Claspin protein is predicted to be intrinsically disordered. The switch on and off of the DNA and protein interaction activities of Claspin through phosphorylation suggests a possibility that intrinsically disordered protein regions may undergo structural transition upon phosphorylation or ligand interaction. Trypsin sensitivity suggests the structural changes are induced by phosphorylation. We have predicted the probability of IDR and structured domains in Claspin derivatives in which all the serine/ threonine residues are converted to aspartic acid/ glutamic acid (ST/DE mimicking the phosphorylated form) or alanine (ST/AA mimicking the unphosphorylated state). ST/DE is predicted to be highly disordered over the entire polypeptide as is the wild-type. On the other hand, the disordered probability significantly decreases in ST/AA over the entire polypeptide compared to the wild-type. Furthermore, 6 globular domains (GD1[41-141aa], GD2[291-341aa], GD3[491-543aa], GD4[711-955aa], GD5[1102-1137aa], and GD6[1194-1339aa]) were predicted in ST/AA (**Supplementary Fig. 3**). There results support the idea that disordered segments are converted to structured domains to enable DNA binding and protein interactions. Trypsin sensitivity correlates well with the disordered structure and the absence of structured domains (required for DNA binding or protein interaction). The disordered form of Claspin is much more sensitive to Trypsin digestion compared to the protein with structures.

### Predicted structures and functions of Claspin during cell cycle

During G1 phase, Claspin adopts a closed conformation, in which the C-terminal segment (AP) and N-terminal segment (DNA binding domain) interact with each other. This interaction suppresses its DNA binding but maintains its ability of helicase inhibition. At the G1-S boundary, Cdc7, CDK or another kinase is recruited to AP on Claspin, phosphorylating more predominantly the C-terminal segment. This is suggested by the finding that the C-terminal polypeptide is more efficiently phosphorylated on the N-C polypeptide complex (**Fig. 5B and C**). This could facilitate the conversion of the closed form to an open conformation by destabilizing the N-C interaction. We also cannot rule out the possibility that a phosphatase removes phosphates from Claspin, activating the DNA binding and protein binding. After Calyculin A treatment (or during the late S-G2-M phase), Claspin would be hyperphosphorylated, resulting in loss of DNA binding and protein interaction and impaired inhibition of helicase. On the other hand, complete removal of phosphorylation in vitro by λPPase leads to Claspin with a more rigid structure (resistant to trypsin) (**Fig. 2C**), which is proficient in DNA binding, protein interaction and helicase inhibition. Phosphorylation in vitro of dephosphorylated Claspin by Cdc7 does not impair its DNA binding activity (**Fig. 1C**), consistent with its expected role as an activator of Claspin. A hyperphosphorylated form of Claspin, generated by Calyculin A treatment or potentially in late S/G2 phase, is deficient in all the activities.

The findings in this communication not only lend a clue to solving the undissolved issue of Claspin-mediated replication regulation by showing that phosphorylation changes the conformation as well as biochemical activities of Claspin, but also provide an important insight into a general issue of biological importance, namely how disordered proteins interact with other molecules and operate during various biological reactions. Future studies will be needed to clarify exactly how cell cycle-dependent phosphorylation/ dephophosphorylation changes the local and overall conformation of Claspin and activates initiation and facilitates the fork progression, and how the replication fork stress changes the IDR state of the protein and leads to activation of Chk1. The availability of the Claspin mutants predicted to form globular domain structures (ΔAP and AP[DE/A] mutants and also see **Supplementary Fig. 3**) will provide an excellent opportunity to determine the defined structures of Claspin important for interaction with DNA and other replication factors.

## Methods

### Cell lines and Enzymes

293T, U2OS and HCT116 were obtained from ATCC. Kinase proteins Cdc7 (05-109), CK1γ1 (03-105), and Chk1 (02-117) were purchased form Carna Biosciences. CK2 (P6010) and λPPase were purchased form NEB Japan Inc. Calyculin A was purchased form Cell Signaling Technology, Inc.

### Plasmid construction

The Claspin-encoding DNA fragment (XhoI/XbaI fragment) of CSII-EF MCS-6His-Claspin-3Flag or CSII-EF MCS-6His-Claspin-HA plasmid DNA was replaced by DNA fragments encoding various portions of Claspin, amplified by PCR, to express truncated forms of Claspin. To express Claspin mutants, ST/A and ST/D, the Claspin coding DNA for 1-350aa, in which all serines/ threonines were replaced with alanine (ST/A) or phospho-mimic aspartic acid (ST/D), were synthesized (Integrated DNA Technologies, Inc.). The ST/A or ST/D fragment and PCR-amplified Claspin (351-1339aa) fragment were inserted into CSII-EF MCS-6His-Claspin-3Flag-P2A-Bsd (Blasticidin) using In-fusion cloning method to construct lentiviral expression vectors for the full-length Claspin ST/A or ST/D mutant.

### Expression of recombinant proteins in 293 T cells

1.6 µg of expression plasmid DNA in 100 µl of 150 mM NaCl were mixed with 100 µl of 150 mM NaCl supplemented with 7 µl of 1 mg/ml PEI (polyethylenimine ‘MAX’ Polyscience, Inc.). After 30 min incubation at a room temperature, the solution was added to 293 T cells cultured in six well plates with 2 ml of fresh DMEM in each well^46^.

### Protein purification

293T cells (15 cm dish, 3–5 plates) were incubated for 40 h rafter transfection, harvested and lysed as described before^10^. The proteins bound to anti-Flag M2 affinity beads (Sigma-Aldrich) were recovered from the supernatants and washed by Flag wash buffer (50 mM NaPi (pH 7.5), 10 % glycerol, 300 mM NaCl, 0.2 mM PMSF and PI tablet), and bound proteins were eluted with Flag elution buffer [50 mM NaPi (pH 7.5), 10 % glycerol, 100 mM NaCl, 200 µg/ml 3xFlag peptide (SIGMA), 0.1 mM PMSF and PI tablet]^46^.

### Immunoprecipitation

293T cells transfected with the indicated Claspin-expressing plasmid were treated with 10 µM RO3306 (RO) and 20 µM Roscovitine ® for 2 hr, or 100 nM Calyculin A (CA) for 30 min before cell harvest. Cells were lysed with native lysis buffer (40 mM Hepes-Na[pH7.5], 130 mM Na-glutamate, 1 mM Mg-acetate, 0.1 mM ATP, 0.5 % Triton X-100), and a half of non-treated cell lysate were incubated with λPPase in the presence of 1 mM MnCl_2_ at 25°C for 30 min. Cell lysates were incubated with anti-Flag-M2 antibody beads at 4°C for 50 min, and the beads were washed four times with lysis buffer. Bound proteins were eluted from beads by boiling in 1xSDS sample buffer, and samples were analyzed by immunoblotting using anti-HA, anti-Flag, and anti-PCNA antibodies.

### Proteolytic digestion with trypsin

Purified Claspin (10 ng/µl) treated with λPPase or that purified from Calyculin A-treated cells was subjected to proteolytic digestion with 0.01 ng/µl trypsin (Roche, sequencing grade) for 0, 2, 5, 10, 15, 20, 30 and 60 min, or with various concentrations of trypsin (0.025, 0.05, and 0.1ng/µl trypsin) for 5 min in 25 mM Tris-HCl (pH 7.9), 50 mM KCl, 1 mM DTT and 0.1% Tween 20. Samples were incubated around 25°C and 12 µl samples were collected and quenched with SDS-sample buffer. Samples were boiled and loaded onto an SDS– PAGE gel. Gels were stained with Coomassie blue and band intensities of protein were measured using Multi Gauge software.

### Preparation of labeled Y-fork

^32^P-end-labeled Y-fork substrates were prepared by annealing oligonucleotide dT_20_-25R with an unlabeled 60mer-dT_50_, as indicated previously^47^. To prepare IR_800_-labelled Y-fork substrates, 5’-end-amino-tethered primer 37mer-dT_25_ (5’-GTTTTCCCAGTCACGACGTTGTAAAACGACGGCCAGT-dT_25_-3’) was labeled with IRDye800CW infrared dyes (IR_800_) and purified by Sep-Pak C18 cartridge chromatography. IR_800_-labelled sense oligonucleotide (37mer-dT_25_, 10 pmol) was mixed with partially complementary antisense oligonucleotide dT_20_-37R (5΄-dT_20_-ACTGGCCGTCGTTTTACAACGTCGTGACTGGGAAAAC-3΄) (12 pmol) in 20 mM Tris–HCl (pH 7.4) and 10 mM MgCl_2_, and incubated at 96° C for 3 min, 65°C for 1 hr, and then 37°C for 10 min.

### DNA binding and DNA helicase assay

DNA helicase and DNA binding activities were examined in reaction mixtures (12 µl) containing 5 mM creatine-phosphate, 20 mM Tris-HCl (pH 7.5), 10 mM Mg-acetate, 1 mM DTT, 0.1 mM EDTA, 20 % glycerol, 50 µg/ml BSA and 60 mM K-glutamate in the presence of the ^32^P-end-labeled fork DNA substrate. In the DNA helicase assay, 10 mM ATP and unlabeled trap DNA were added and incubated at 37°C for 60 min. After termination of the reaction by adding the stop buffer (final 20 mM EDTA, 0.1 % SDS and 0.25 % BPB), products were separated on 10% acrylamide/1× TBE gel. DNA binding assays were conducted at 30°C for 30 min without ATP. All of samples were run on 5% non-denaturing gel containing 0.5× TBE and 5% glycerol. For DNA binding assays with IR_800_-labeled substrates, samples were run on 5-20% nondenaturing gels containing 1× TBE or 1× Tris-glycine as indicated.

### PLA assay

The proximity ligation assay (PLA) was conducted to determine potential protein–protein interactions using the Duolink in situ detection kit (Sigma-Aldrich) in accordance with the manufacturer’s instructions. Combinations of antibodies from different species were used for detection. Briefly, cells were fixed in 4 % paraformaldehyde in PBS for 10 min and permeabilized in PBS–0.5% Triton X-100 for 10 min. Cells were blocked in blocking buffer (1 % BSA and 0.1 % Tween20 in PBS) for 2 hr to avoid any non-specific binding. The primary antibodies directed against HA (1:1800; MMS-101R; Covance), Flag (1:150; ab245892; Abcam), and FKBP12 (1:150; ab58072; Abcam) were diluted in blocking buffer in PBS overnight at 4°C. Combination of secondary anti-rabbit PLUS and anti-mouse MINUS probes was used followed by hybridization, ligation, and amplification steps. Nuclei were counterstained with Hoechst33342. Red fluorescent spots (proteins located within 40 nm of each other) were visualized with a standard Keyence BZ-X 710 All-in-One fluorescence microscope or Zeiss LSM980 confocal microscope.

### Kinase assay of various Claspin

Purified 300 ng each of wild type (WT), mutants (ΔAP, ST/A, and ST/D), and N-terminal (#2) or C-terminal (#13 and #27) polypeptides of Claspin was treated with 100 unit λPPase for 30 min at 30°C and the reaction was stopped by adding phosSTOP at 25°C for 1 h. Samples were mixed with 25 ng various kinases (Cdc7, CK1γ1, CK2 or Chk1) and incubated at 30°C for 30 min in kinase buffer (50 mM Tris-HCl[pH7.5], 10 mM MgCl_2_, 0.01% Brij35, 0.1 mM EDTA, and 2 mM DTT) with 1.8 µCi [γ-^32^P] ATP and 1 mM ATP. In some experiments, #13 polypeptide was incubated with a first kinase indicated and cold 1 mM ATP in buffer for 30 min at 30°C, followed by incubation with a second kinase with γ-^32^P ATP. After addition of 5x sample buffer, samples were boiled and applied to SDS-PAGE. After Coomassie blue staining, the gel was autoradiographed.

### Generation of HCT116 Claspin-mAID

HCT116 Claspin-mAID was generated as described previously^48^. The guide RNA sequence targeting CLSPN (human Claspin) locus was TGGAGAGCTAACACCATCAA(AGG) [protospacer adjacent motif (PAM) sequence is indicated in parentheses].

### Cell lines for ectopically expressing Claspin

The relevant cell lines were constructed using lentivirus infection, followed by antibiotics selection. The lentiviruses were generated by co-transfection of Uno8-4 vector containing either WT or mutated Claspin (ST/A and ST/D), pCAG-HIVgp, and pCMV-VSV into HEK293FT cell at a molar ration of 1:1:1. The isolated lentiviruses were used to infect mAID (HCT116) cells. Cells were selected with blasticidin (Bsd) at 10 µg/mL.

### Colony formation assay

Colony formation assay was conducted by Giemsa stain solution (079-04391, Fujifilm) by following the manufacturer’s guidelines. Briefly, cells were treated with or without HU for 3 hr. Cells were then trypsinized and 100 cells were seeded into a well of a 6-well plate. After 3 weeks, cells were washed with PBS three times and fixed with methanol for 30 sec. Methanol was then removed and Giemsa stain solution were added to cells for 30 min. Cells were then rinsed with ddH_2_O for 15-30 sec and air dried. Colony numbers were counted and presented.

## Acknowledgements

This work was supported by JSPS KAKENHI (Grant-in-Aid for Scientific Research (A) [Grant Numbers 20K21410 and 20H00463 (to H.M.)]; and Hirose international scholarship foundation (to C-C.Y.). We thank Tomohiro Iguchi for his help in making lentiviruses. We thank Masato Kanemaki (National Institute of Genetics) for providing IAA-inducible protein degradation system. We would like to thank all the members of the laboratory for helpful discussion.

## Author contributions

Z. You performed the experiments and contributed to the analysis of the results described in the paper. H-W. Hsiao, and C-C. Yang performed some of the experiments. H. Goto constructed HCT116 Claspin-mAID cell lines. Z. You and H. Masai designed the experiments and wrote the manuscript with input from all authors. H. Masai obtained fundings and has overall responsibility for the study.

