## Supplementary Figures 1-3 for "Phosphorylation inhibits intramolecular interactions, DNA-binding and protein interactions of Claspin through disordered/ structured conformation transition"

### Supplementary information

Supplementary Fig. 1

A.

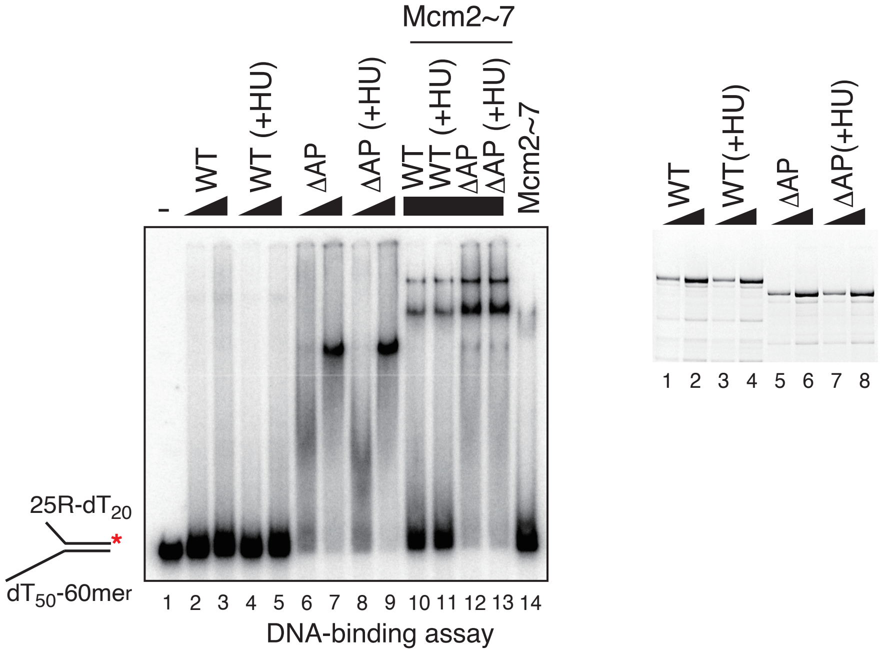


B.

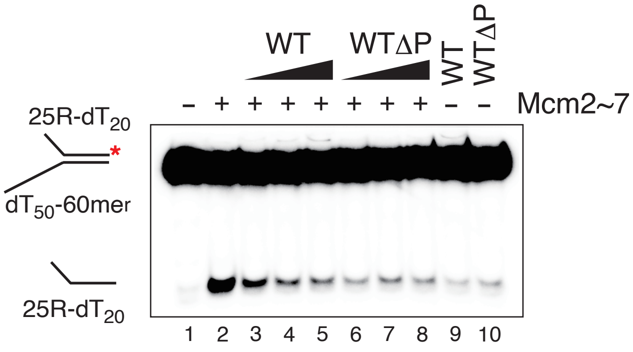


**Supplementary Figure 1. Claspin purified from the HU-treated and non-treated cells equally binds to DNA.**

**A.** Gel shift assays were conducted with $^{32}$P-labeled Y-fork DNA. Proteins indicated were purified from 293T cells, non-treated or treated with 2 mM HU for 1.5 hr (+HU), before cell harvest. Proteins used for assays were as follows; WT, WT(+HU), ΔAP and ΔAP(+HU), 150 ng and 300 ng for each protein in lanes 2-9 and 300 ng for lanes 10-13; Mcm2~7, 75ng. Y-forked DNA, 50 fmole. The protein preparations used for the assays are shown in the right panel (CBB staining).

**B.** Effects of phosphorylated and dephosphorylated Claspin on DNA helicase activity of Mcm2~7. Proteins added; highly purified fraction of Claspin protein (WT and WTΔP) 50, 100 and 200 ng; Mcm2~7, 50ng. $^{32}$P-labeled Y-fork DNA, 15 fmole.

Supplementary Fig. 2

A.

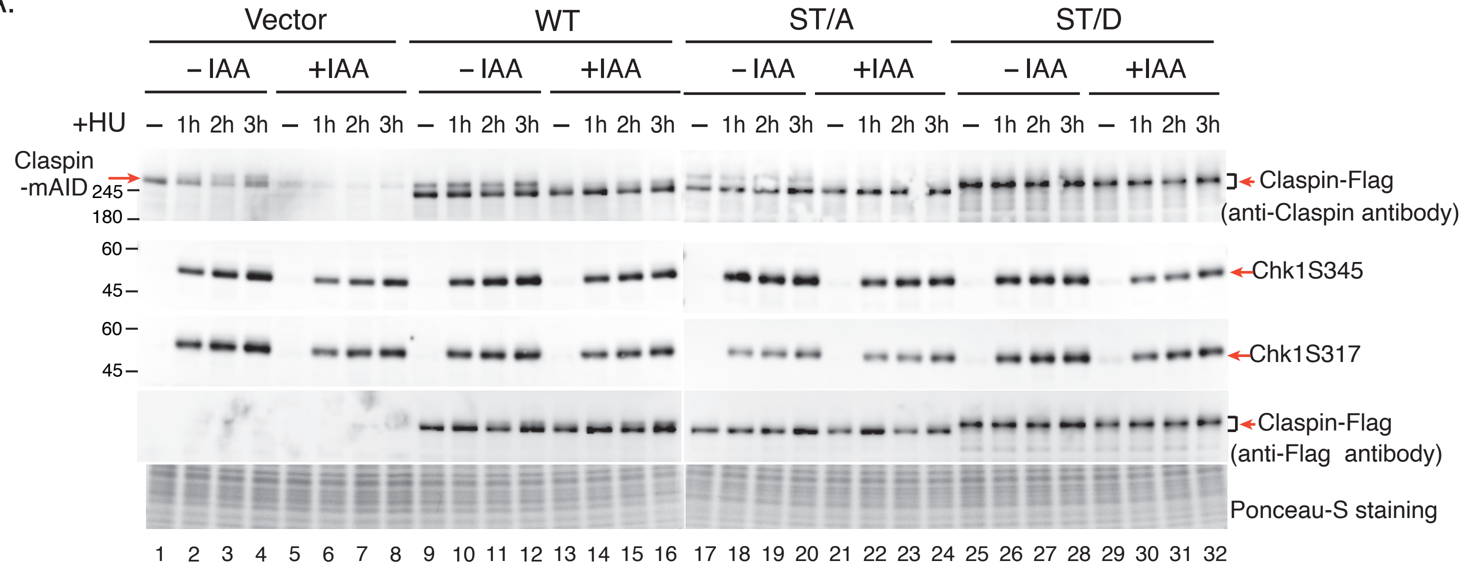


B.

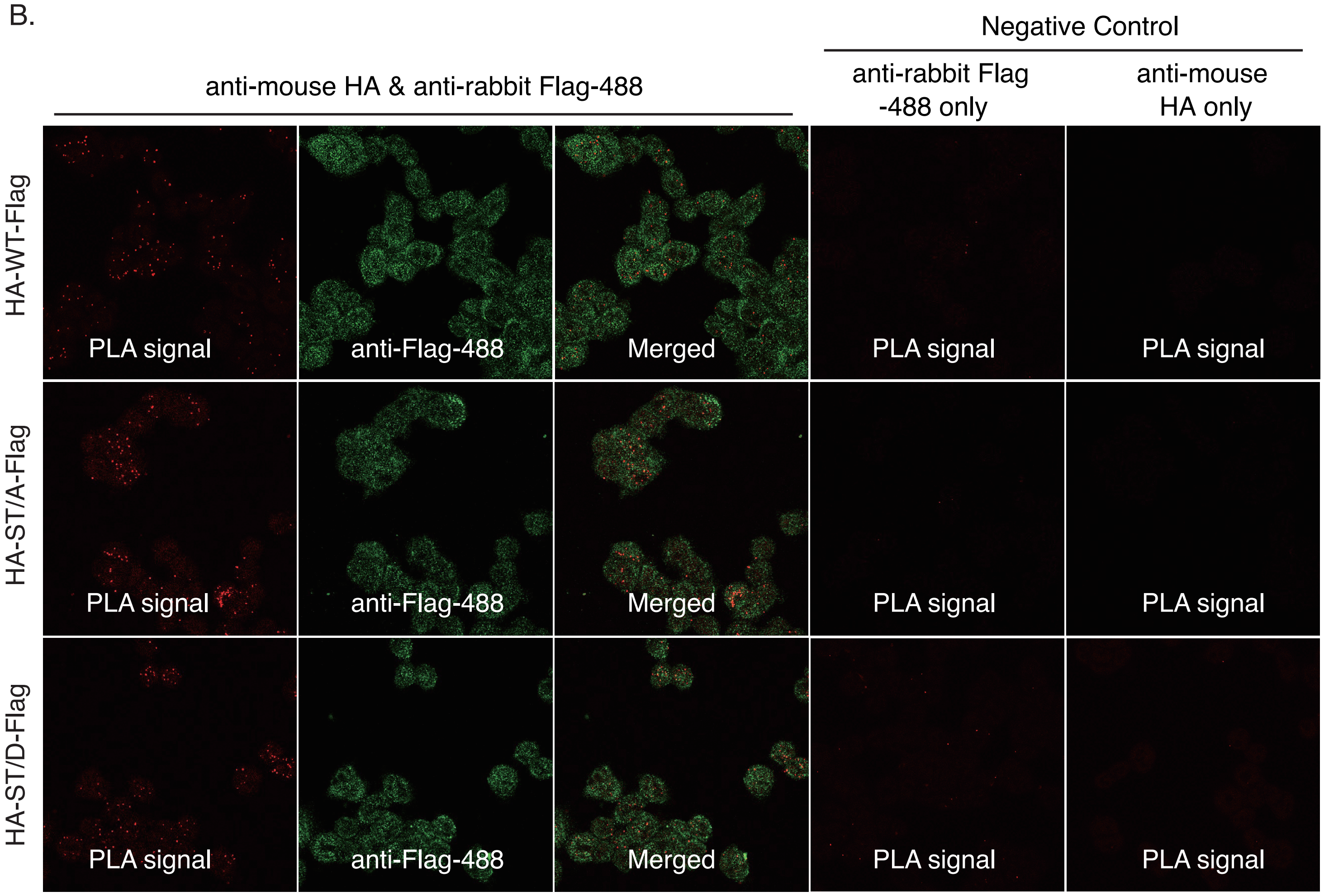

**Supplementary Figure 2. ST/D mutant is defective in replication stress responses**

**A**. HCT116 Claspin-mAID cells expressing wild-type (WT) or mutant forms (ST/A and ST/D) of Claspin were treated with IAA for 7 hr or untreated. Prior to the cell harvest, cells were untreated or treated with 10 mM HU for 1 hr, 2 mM HU for 2 and 3 hr, and were analyzed by western for expression of various proteins as indicated.

**B**. PLA with HCT116 Claspin-mAID cells expressing different forms of HA-Claspin-Flag (WT, ST/A, and ST/D). After treatment with IAA for 4 hr, PLA signals were observed under confocal microscopy using anti-mouse HA and anti-rabbit Flag-488 primary antibodies. The right two columns represent negative controls, PLA signals detected only with a single primary antibody (anti-rabbit Flag-488 primary antibody [$2^{nd}$ column from right] or anti-mouse HA primary antibody [the right-most column])

Supplementary Fig. 3

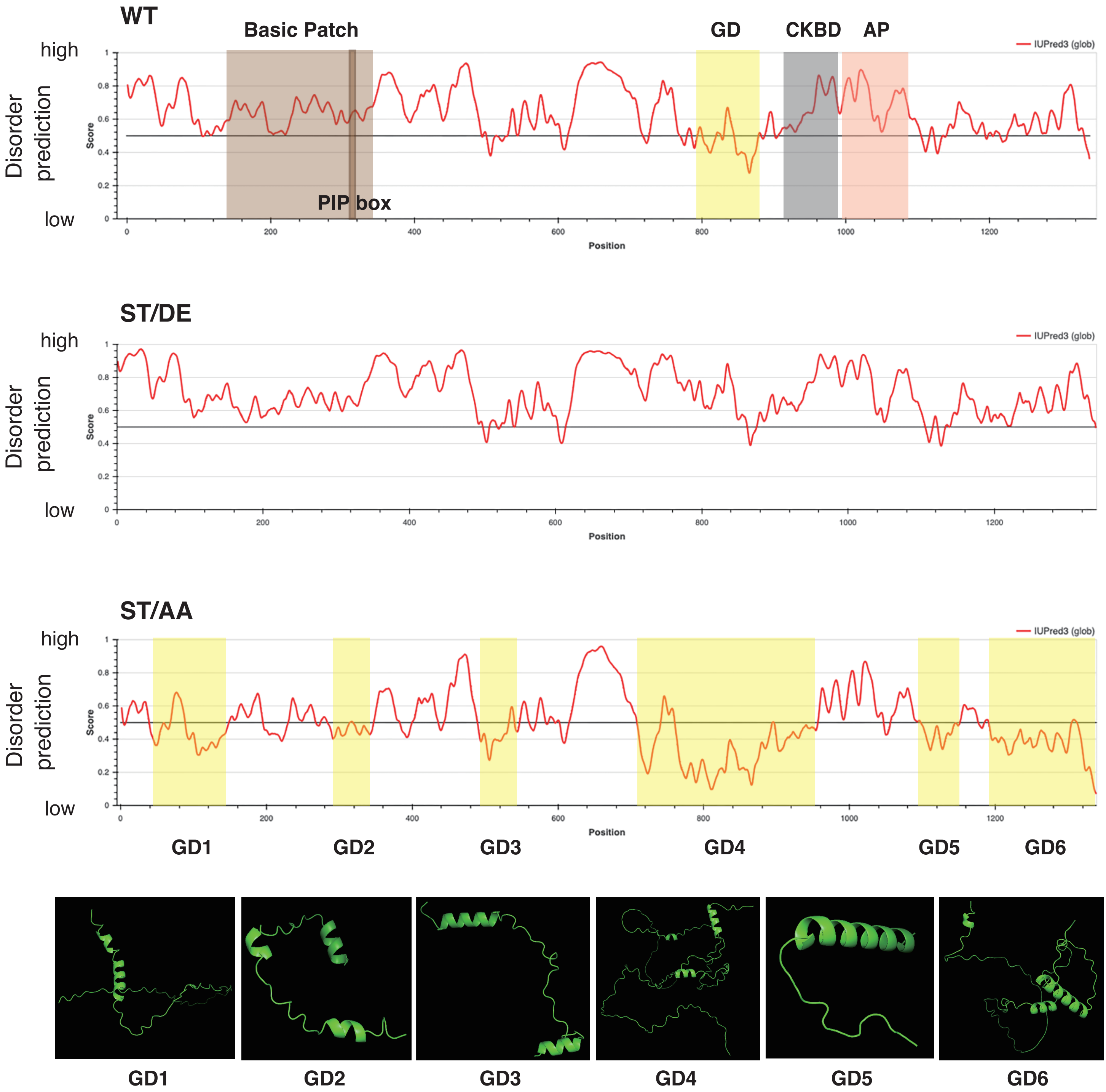


**Supplementary Figure 3. Prediction of disordered and structured segements in Claspin and its phosphomimic and phosphodefective derivatives.**

The wild-type Claspin (WT) or its derivatives ST/AA or ST/DE in which all the serine and threonine residues were replaced, respectively, by alanine or by aspartic acid (for serine) and glutamic acid (for threonine) were predicted by "IUPred3 structural domains" (https://iupred.elte.hu). The segments highlighted in yellow boxes represent predicted globular domains (GD1[41-141aa], GD2[291-341aa], GD3 [491-543aa], GD4[711-955aa], GD5[1102-1137aa], and GD6[1194-1339aa] in ST/AA and GD [791-876aa] in the wild-type). Boxes with other colors indicate the signatures of Claspin molecules as shown. Y-axis shows the prediction score for IDP. The structures of GD1-6 predicted by AlphaFold 2 are shown at the bottom of the figure.
